# A zero percent plastic ingestion rate by silver hake (*Merluccius bilinearis*) from the south coast of Newfoundland, Canada

**DOI:** 10.1101/301630

**Authors:** France Liboiron, Justine Ammendolia, Jacquelyn Saturno, Jessica Melvin, Alex Zahara, Natalie Richárd, Max Liboiron

**Author notes:** Corresponding author Address: SN1107A, Memorial University of Newfoundland, 230 Elizabeth Avenue, St. John’s, NL, A1C 5S7. **Declaration of Interest** None. **Terms:** - Frequency of occurrence: the number of individual fish within a sample population that have ingested plastics, regardless of how many plastics they ingested (%FO).

## Abstract

Silver hake, (*Merluccius bilinearis*), contributes significant biomass to Northwest Atlantic ecosystems. The incidence of plastic ingestion for 134 individuals collected from Newfoundland, Canada was examined through visual examination of gastrointestinal contents and Raman spectrometry. We found a frequency of occurrence of ingestion of 0%. Through a comprehensive literature review of globally published fish ingestion studies, we found our value to be consistent with 41% (*n*=100) of all reported fish ingestion rates. We could not statistically compare silver hake results to other species due to low sample sizes in other studies (less than *n*=20) and a lack of standardized sampling methods. We recommend that further studies should 1) continue to report 0% plastic ingestion rates and 2) should describe location and species-specific traits that may contribute to 0% ingestion rates, particularly in locations where fish consumption has cultural and economic significance.

## Introduction

The province of Newfoundland and Labrador, Canada plays a critical role in the nation’s fishing industry. Despite the collapse of the Atlantic cod (*Gadus morhua*) stock and the province’s subsequent moratorium on Atlantic cod in 1992, the fishing industry still contributes 3.1 billion dollars in revenue and employment of over 17,000 individuals in the remote province (Bavington, 2010; Campling et al., 2012; Governement of Newfoundland and Labrador, 2016). One of the ways the Newfoundland fishery has survived the cod collapse is by diversifying what is fished (Bavington, 2010). Although the slender schooling fish, silver hake (*Merluccius bilinearis*), makes up a significant biomass of the Northwest Atlantic ecosystem (where biomass is a measurement used by the fisheries to describe the total mass of organisms in a given area), there is no established silver hake fishery in Newfoundland (Bayse and He, 2017; Garrison and Link, 2000), making it a potential species through which to further diversify the fishery in the province. Silver hake have a broad geographic distribution in the Atlantic Ocean ranging from South Carolina, U.S.A. to Newfoundland, Canada (Bayse et al., 2016; Bigelow and Schroeder, 1953). Since the 1950s, there have been commercial fisheries in North America for various species of hake (Helser and Alade, 2012; Pitcher and Alheit, 1995), and the Canadian and American silver hake fishery produces approximately 8-9,000 metric tons/yr and 6-7,000 metric tons/yr, respectively (DFO, 2015; NMFS, 2017). Despite the size of the silver hake fishery in the North Atlantic, Canada only became involved in the fishery in 2004, and Canadian fleets fish exclusively in the Scotian shelf south of Newfoundland (DFO, 2015). The catch from the American fleets mainly comes from the New England fishery fished in the Gulf of Maine and Georges Banks region, where the populations are relatively stable with no scientific evidence of past overfishing (Bayse and He, 2017; Morse et al., 1999; New England Fisheries Management Council, 2012). Silver hake is an economically important fish species as its flesh has been found to be an alternative fish product for the manufacture of surimi, a seafood analog (e.g. imitation crab meat), that is consumed worldwide (Lanier, 1984).

Fragmented or manufactured plastics are ubiquitous in marine environments. Plastics that are small in size (< 5 mm) are known as microplastics, a size class which accounts for more than 90% of all marine plastic particles (Eriksen et al., 2014). The abundance and small size of marine microplastics makes them highly bioavailable for ingestion by marine animals. It has been reported that plankton - some of the smallest marine animals which form the base of the marine food web - can successfully ingest microplastics (Cole et al., 2014). This is especially concerning given that plastics contain contaminants enmeshed during manufacture such as plasticizers, colourants, and flame retardants (Colton et al., 1974; Lithner et al., 2012) as well contaminants that accumulate on plastics from the surrounding seawater, such as flame retardants like polychlorinated biphenyls (PCBs) and insecticides such as dichlorodiphenyltrichloroethane (DDT), among many other chemicals (Mato et al., 2001; Newman et al., 2015). There is potential for these toxicants to bioaccumulate in marine animals that ingest plastics (Rochman, 2013). The biomagnification of these contaminants into higher trophic levels of the food web (including human consumers), while of great concern, is not fully understood and is an increasing area of study (Bakir et al., 2016; Engler, 2012). Most plastic ingestion studies, particularly in Canada, have focused on seabirds (Bond, 2016; Bond et al., 2012; Bond and Lavers, 2013; Holland et al., 2016; Provencher et al., 2014), while fewer have focused on fish, and only a handful have studied fish destined for human consumption (Choy and Drazen, 2013; M. Liboiron et al., 2016; Rochman et al., 2015).

The vulnerability of marine animals to the ingestion of plastics will depend on their ecological niche. For example, given that plastics exist at various depths but accumulate in pelagic and benthic environments, fish that occupy bathymetric ranges that correspond to high marine plastic abundance may be more susceptible to plastic ingestion. For instance, Neves et al., (2015) found that pelagic species consumed more plastic particles when compared to the sampled benthic species from off the Portuguese coast. This is consistent with reports of high concentrations of marine plastics in the ocean’s surface layer (Cózar et al., 2014; Eriksen et al., 2014). To date, the relationship between fish species’ bathymetric depth range and plastic ingestion have not been regularly or systematically examined within or across published studies.

Silver hake are a species of fish we hypothesize are likely to ingest plastics. They are a demersal species, found in depths ranging from 55-914 m (Lloris and Matallanas, 2005), where they feed from both surface and benthic environments. Because they are predators, silver hake are assumed to be a species likely to ingest food that contains plastics (via secondary ingestion) in environments that contain plastics (particularly surface waters where plastics accumulate) (Andrady, 2003; Eriksson and Burton, 2003; Romeo et al., 2015). Between the ages of 1-3 years, silver hake opportunistically feed on invertebrates, transitioning to a more piscivorous diet at maturity (Vinogradov, 1984; Waldron, 1992), potentially increasing the size of plastics they may ingest via secondary ingestion. In fact, individual silver hake > 40 cm typically feed exclusively on fish (Durbin et al., 1983; Langton, 1982), and as adults commonly exhibit cannibalistic behavior (e.g. diets consist of ~10% frequency of occurrence of juvenile conspecifics) (Bowman, 1983; Bowman, 1975). Variation from this piscivorous diet can arise as a result of seasonality, combined with the opportunistic nature of predation (Waldron, 1992). For instance, during the spring adults mostly consume fish, while diverse prey items like crustaceans and molluscs may also be preyed upon during the summer (Waldron, 1992). Generally, however, silver hake feed primarily on pelagic species that in turn feed from the surface of the water where plastics tend to accumulate. In relation to other fish species in the region (e.g. cod and haddock), silver hake are more selective in their feeding habits and the species of prey consumed are less diverse (Bowman, 1975). Although the feeding ecology of this fish has been well-studied, a search of published English-language scientific literature has returned no research examining plastic ingestion by silver hake, despite feeding habits that position them to do so.

The silver hake in this study were collected from an area of suspected high plastic pollution, off the south coast of Newfoundland, both within the Gulf of St. Lawrence and just outside of it, along the southwestern Grand Banks (Fig. 1). This sample area is expected to have a higher representation of plastic pollution compared to more northern Newfoundland waters fed by the Labrador current (Liboiron et al., 2016). The Gulf of St. Lawrence is surrounded and enclosed by five Canadian provinces, is home to intense fishing (especially in the case of the Grand Banks) and shipping activity, and sits at the mouth of a river draining a large portion of mainland Canada and the United States (Fisheries and Oceans Statistical Services, 2016; The St. Lawrence Seaway Management Corporation, 2016). The numerous pathways of introduction for marine plastics in the Gulf of St. Lawrence and the high fishing intensity on the Grand Banks makes this sample site ideal for the investigation of plastic ingestion. Regardless of the effect that the feeding behavior of silver hake may have on their ingestion of plastic, silver hake of this region are expected to be at a higher risk of plastic ingestion than in other regions around Newfoundland. If %FO of plastic remains low despite environmental plastic concentrations that are expected to be high, silver hake may represent a safe new fishery option within the context of plastics in fish in the face of increasing plastic loads in the future.

**Fig. 1.**
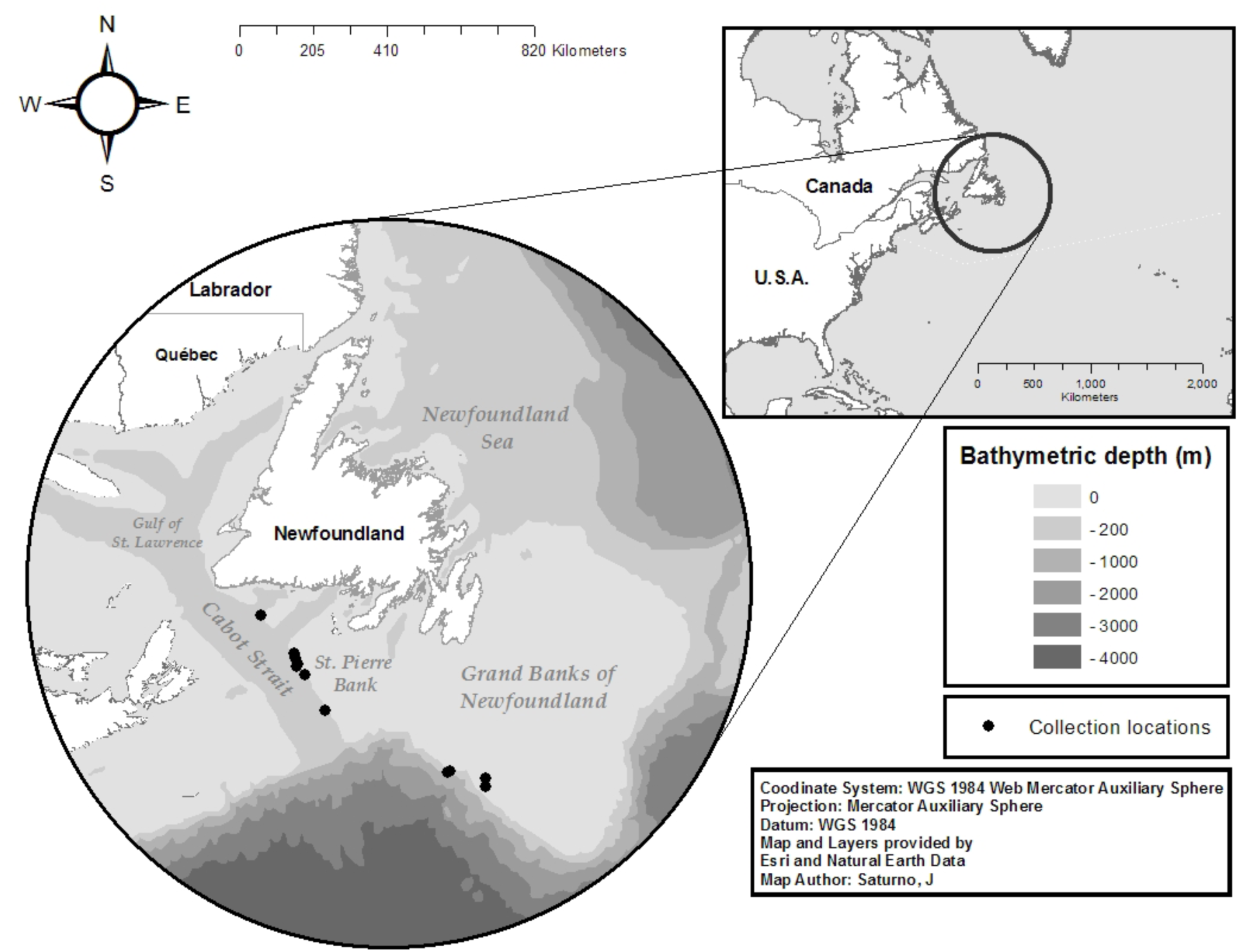
Collection locations sampled for silver hake off the southern coast of Newfoundland.

## Methods

### Collection of silver hake

Individual silver hake were collected by trawling from the RV Celtic Explorer research ship, from April 27 to May 6, 2016 by Fisheries and Oceans Canada and the Fisheries and Marine Institute of Memorial University of Newfoundland (Table 1; Fig. 1). Of the 175 silver hake collected off of the south coast of Newfoundland, 41 individuals were eliminated from analysis due to compromised gastrointestinal tracts (inverted, split or detached stomachs occurring during the course of trawling or initial processing), resulting in a total of 134 fish for this study. Body length and sex of each individual was recorded, and entire gastrointestinal (GI) tracts were were removed, individually bagged and tagged, and frozen aboard the RV Celtic Explorer for later transport to the laboratory.

**Table 1.** Set details for the silver hake samples collected in the Northeast Atlantic used in this analysis.

| Location | No. of fish collected | Tow # | Depth (m) | Date | Latitude (°North) | Longitude (°West) | Duration of tow (min) |
| --- | --- | --- | --- | --- | --- | --- | --- |
| Burgeo Bank | 9 | 1 | 198 | April 27, 2016 | 47.1063 | -58.1531 | 15 |
| St. Pierre Bank | 85 | 5 | 309 | April 28, 2016 | 46.4600 | -57.3171 | 17 |
|  |  | 7 | 231 |  | 46.3620 | -57.3019 | 16 |
|  |  | 9 | 328 | April 29, 2016 | 46.2674 | -57.2414 | 15 |
|  |  | 10 | 307 |  | 46.2339 | -57.2574 | 15 |
|  |  | 12 | 144 | April 30, 2016 | 46.0900 | -57.0601 | 15 |
|  |  | 15 | 344 | May 1, 2016 | 45.4672 | -56.5412 | 17 |
| Southern Grand Bank | 40 | 31 | 218 | May 5, 2016 | 44.3578 | -53.4863 | 15 |
|  |  | 32 | 118 |  | 44.3885 | -53.4362 | 16 |
|  |  | 35 | 124 | May 6, 2016 | 44.2549 | -52.5302 | 16 |
|  |  | 36 | 220 |  | 44.1087 | -52.5353 | 16 |

### Laboratory procedures

#### Visual Analysis

We undertook precautions to avoid cross-contamination of plastics, which included: rinsing or wiping down all tools with water and Kimwipes, including the microscope lens and plate, Petri dishes, and sieves before use; hands were washed; cotton lab coats worn; and hair was tied back. After each dissection, we closely examined our hands and tools for any microplastics that may have adhered. We kept a control dish to collect microplastics, specifically microfibers, that originated within the lab, the contents of which were then used to compare to any microfibers found in the fish for exclusion purposes. However, microfiber contamination precautions were not taken during the collection period onboard the ship.

This study followed methods used by Liboiron et al. (2016) (which were in turn adapted from van Franeker (2011) and Avery-Gomm (2016)) to allow for comparability across studies done in the region of Newfoundland. The bagged GI tracts were thawed in cold water for approximately 30 min prior to dissection. A double sieve method was used by placing a 4.75 mm (#4) mesh stainless steel sieve directly above a 1 mm (#18) mesh stainless steel sieve. The 4.75 mm sieve served to separate mesoplastics and larger GI contents from microplastics and smaller items. The lower threshold of 1 mm was selected as anything smaller cannot be reliably identified with the naked eye (Song et al., 2015). GI tracts were placed in the 4.75 mm sieve and an incision was made using fine scissors, running from the esophagus, to the stomach, through the intestines, then to the anus. The contents of the GI tract were gently rinsed with tap water into the sieve to remove contents. The GI lining was closely examined for any embedded plastics and the contents of the sieves were visually inspected. Any identifiable food was recorded, and suspected anthropogenic debris was removed with tweezers and transferred to a Petri dish for later observation under a microscope. Stomach contents were described as containing either no food, small amounts of food, or large amounts of food. “Small amounts” of food described food detritus, wherein the presence of food was not apparent before cutting into the stomach and included items such as shrimp shells, fish scales, fish bones, and small shrimp. “Large amounts of food” described stomach contents that created bulk in the GI tract identifiable before cutting into it, and in many cases contents expanded the stomachs or even protruded from their stomachs into the esophagus.

Items suspected to be anthropogenic debris were placed into a paper filter, labeled with the fish ID, and left to dry. Once dried, they were examined with a dissecting microscope (Olympus SZ61, model SZ2-ILST, with a magnification range of 0.5–12×) with both reflected oblique and transmitted light and, if necessary, under a compound microscope (Ecoline by Motic, Eco Series, with a magnification range of 4-100×).

#### Raman spectrometry

To validate our visual inspection methods, six particles initially identified as non-plastic were prepared for Raman micro-spectrometry. This analysis was selected because it has been proven to be a nondestructive spectroscopic method that can accurately differentiate between nonorganic and organic particles. Analyses can be used to identify the molecular structure of a particle by characterizing the vibrational properties of the sample (Imhof et al., 2012; Lenz et al., 2015). The high resolution of this method can in many cases successfully identify the specific polymer of a tested plastic item (Lenz et al., 2015).

Particles were rinsed in ethanol and allowed to dry. Samples were put onto a silica wafer with a known Raman spectrum of 520 cm^-1^ characteristic peak. Results were analyzed by using WiRE 3.4 software that was connected to the Raman micro-spectrometer (Reinshaw InVia with 830 nm excitation) at an objective of 20x. In order to limit fluorescence and prevent samples from being burnt, the laser power did not exceed 5%. A reference spectra was used to ensure that the Raman spectrum for each sample could be compared to the following common marine plastic polymers: acrylonitrile butadiene styrene (ABS), cellulose acetate, polyamide (PA), polycarbonate (PC), polyethylene (PE), polyethylene terephthalate (PET), poly(methyl methacrylate) (PMMA), polypropylene (PP), polystyrene (PS), polyurethane (PU) and polyvinylchloride (PVC) (Bråte et al., 2016; Engler, 2012; Lenz et al., 2015; PlasticsEurope, 2016).

#### Comprehensive literature review

In addition to laboratory analysis, we conducted a systematic literature review of published English-language ingestion rates of all species of fish, globally, in order to support our hypothesis that although all species of fish - regardless of their bathymetric range - are likely to ingest plastics, certain bathymetric zones present a greater risk in terms of the presence and ingestion of plastics. We searched for English-language papers within JSTOR and Web of Science databases using the following terms in the title, keywords, and abstract: “*microplastic*”; *“plastic*”; “*fish*” and “*ingestion*”. No laboratory studies were included, but both wild or farmed fish were included.

We obtained 22 published studies and disaggregated their ingestion data by species, which provided 242 individual ingestion rates for 207 different species in 64 families (Table S1 in Supplementary Materials). Sample sizes of specific species within examined studies ranged from 1 to 741 individuals, and were often aggregated to produce an overall ingestion rate. We disaggregated these studies into their constituient species and their ingestion rates. Life stages for individuals recorded were either at juvenile or adult stages. Studies that did not provide species-specific plastic ingestion rates are included in Table S1 but not used in Fig. 2–3. In addition to species and ingestion rates, we recorded a study’s geographical location, the family of the species, and whether the samples were freshwater or marine. These data were used to categorize bathymetric range.

**Fig. 2.**
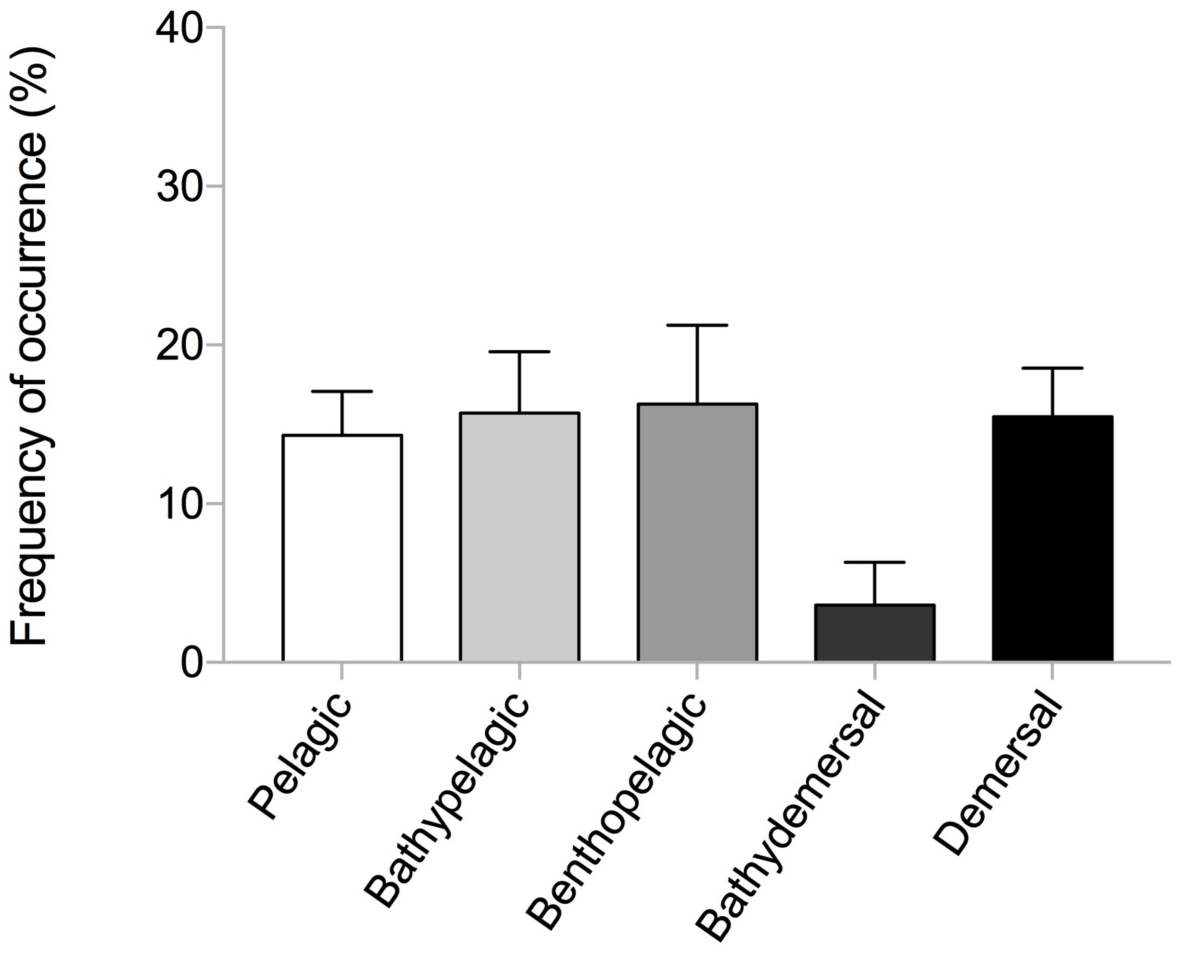
Average frequency of occurrence (%) for species (*n*=7-62) from various bathymetric ranges. Bars represent standard error.

**Fig. 3.**
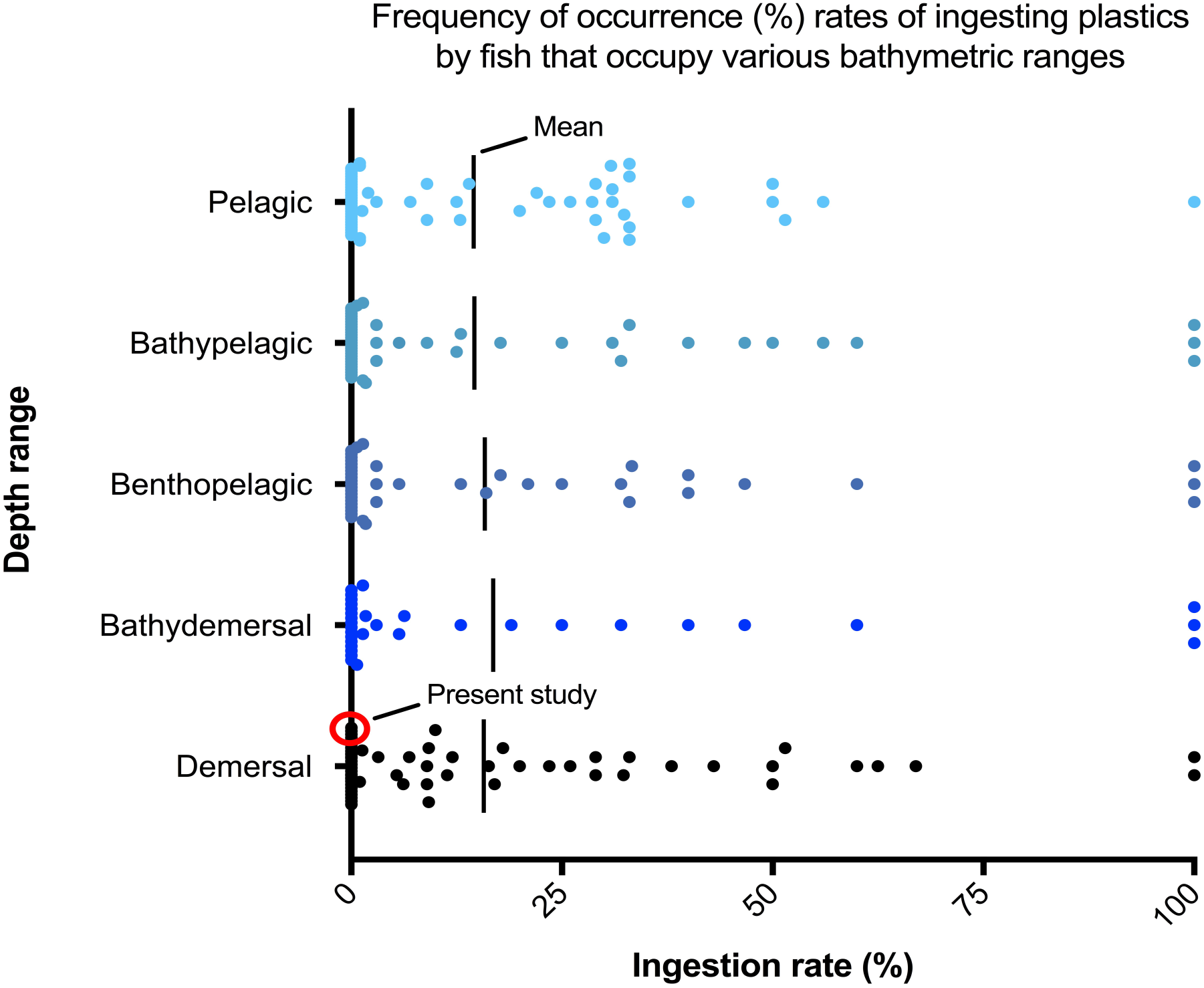
Frequency of occurrence (%) rates of ingesting plastics by fish that occupy various bathymetric ranges (pelagic, bathypelagic, benthopelagic, bathydemersal, demersal), arranged by depth and rate. The demersal silver hake ingestion rate was 0%, as indicated by the red circle. The lines represent the mean for each bathymetric range.

### Data Analysis

#### Silver hake

No data analysis was conducted on plastic ingestion rates as no plastics were recovered. Body length and sex of individuals were analyzed to better characterize life stage and corresponding feeding habits. Length data was summarized using descriptive statistics; including median and mean ± standard deviation. A binary logistic regression was conducted in R Studio version 1.0.143 to test for a relationship between fish length and prey items. All graphs were generated using Prism GraphPad version 7.0 (Prism, GraphPad, 2016) and R Studio version 1.0.143 using ggplot2 (RStudio Team, 2016). Maps were made with ArcGIS version 10.2.2 (ESRI, 2014).

#### Comprehensive literature review

Depth distribution of individual species was determined by using Fishbase (http://www.fishbase.ca) (Froese and Pauly, 2017). Depth range was defined as the minimum and maximum depth of occurrence (m), giving rise to the following categories: pelagic (surface to 200m depth of open water column); bathypelagic (open water column, greater than 200 m in depth); benthopelagic (inhabiting both the seafloor and pelagic waters); demersal (on or near the seafloor, up to 200m depth); bathydemersal (on or near the seafloor, greater than 200 m depth). These five categories were used in Cheung et al., (2007) and are further defined on Fishbase. This database has been used in previous microplastic ingestion studies (Neves et al., 2015; Phillips and Bonner, 2015).

This data was combined with the %FO rates of plastic between fish to see if ingestion rates were impacted by bathymetric range. Using R Studio version 1.0.143, the one-way analysis of variance (ANOVA) was employed to detect significant differences in %FO between different bathymetric ranges. Where the null hypothesis was rejected and significance was determined to be true at the 95% confidence interval, the Tukey honest significant difference post hoc test was conducted to identify which groups contributed to the significant result. In another analysis, “pelagic” and “bathypelagic” categories were combined, as were “bathydemersal” and “demersal” categories for a comparison of relative locations in the water column. We combined both the pelagic and bathypelagic zones because fish from these regions feed in the open water column, while demersal and bathydemersal zones were combined as fish both inhabit and feed on the seafloor. The difference between demersal and bathydemersal fish is that the first inhabits the relatively shallow seafloor, and the second can be found where the seafloor reaches greater depths (similarly, bathypelagic fish inhabit the open water column at greater depths than pelagic fish). These categories better describe fish species based on their relative feeding zones and habits.

### Results

#### Specimen dissections

Of the 134 individual silver hake examined, none were found to contain plastics (0% FO). The 0% plastic ingestion rate is not a result of a lack of feeding by the silver hake collected in this study, as only 9% (*n*=12) of fish analyzed for plastics had stomachs completely devoid of food. The most common food item was shrimp, the remnants (partially digested whole shrimp, or shrimp carapaces) of which were found in 91% of fed fish (*n*=111). The remnants of fish (including partially digested whole fish, bones, otoliths and/or scales) were found in 56% (*n*=68) of fed silver hake individuals. These fish prey items were in all cases too degraded for subsequent dissection as GI tracts were not intact, and the stomach contents of the silver hake were assumed to include any plastics that may have been ingested via secondary ingestion. The prey observed were generally large, and in one instance an ingested shrimp was twice the size of the stomach itself. Smaller food items included arthropods and copepods. Only one individual (<1%) had ingested a strictly benthic invertebrate (crustacean) and there was no evidence of the ingestion of non-biological materials (e.g. sediment). Based on the ingested species found, we can speculate that the silver hake were feeding predominantly in pelagic regions as the vast majority of prey were non-benthic organisms. The lack of non-organic items, including anthropogenic debris, would also suggest feeding above the benthos, as fish that feed on benthic organisms have a tendency to ingest items from the sediment.

Sex and length data was recorded for all fish collected (*n*=175), regardless of the condition of their gastrointestinal tracts. Only 3 individuals (1.7%) exceeded 40 cm in length, the size at which silver hake are expected to be exclusive piscivores. The average length of fish collected was 32 ± 3.5 cm and the most common length (median) was 32 cm. The majority of individuals were between 30 and 40 cm in length (*n*=140; 80%) with only 27 individuals (15%) below this size. Silver hake body length was determined to have no significant relationship with the ingestion of fish versus other prey species (p=0.401). Of the reduced sample size (*n*=134), 71.6% of fish were female (*n*=96), 27.6% were male (*n*=37), and one individual remained unidentified.

#### Raman spectrometry analyses

We analyzed a total of six particles using Raman micro-spectrometry (Fig. S1 in Supplementary Materials). Of these six, one particle, which appeared to be a hair, was not analyzed, as it was too fine to be processed by the machine (Fig. S1F). The remaining five particles did not show any similarity with the reference spectra for plastic polymers. Three of these particles showed identical vibrational characteristics which matched or showed strong similarities to common phases of calcium carbonate, calcite, and aragonite (De La Pierre et al., 2014). The result of this analyses in combination with the visual appearance of the particles, suggests that the particles were bone fragments.

#### Literature review

Multi-species studies are common in the plastic ingestion literature (*n*=17). The 22 studies reviewed here covered a combined species count of 207 species of fish, yielding 242 plastic ingestion rates (%FO) (Table S1). Half of these 22 studies (*n*=11) reported at least one ingestion rate of 0%. Within the 11 studies, there were a total of 100 individual cases of a species with a 0% ingestion rate. These %FO reports spanned 95 different species and made up 41% of all ingestion rates reported in the literature. Most of the plastic ingestion rates reported were associated with relatively low sample sizes, with 67% (*n*=161) comprising a sample size of less than 20 fish. Of these, 81% (*n*=131) were based on sample sizes of less than 10 fish. Studies used different size ranges for identified ingested plastics, which will result in different ingestion rates This is important to keep in mind when comparing overall rates. See table S1 for full details and references.

##### The effect of bathymetric range on ingestion rate

Fish species that had ingested plastics and those that had not inhabit all bathymetric regions. When separated based on bathymetric range, %FO did not differ significantly between pelagic, bathypelagic, benthopelagic, bathydemersal or demersal zones (p=0.782, df=4, *n*=202; Fig. 2–3). The benthopelagic category boasted the highest average %FO (16.3%; *n*=36, SD=29.8) (via Possatto et al., 2011; Anastasopoulou et al., 2013; Neves et al., 2015; Phillips & Bonner, 2016; Rochman et al., 2015; Rummel et al., 2016), followed by the bathypelagic (15.7%; *n*=44, SD=25.6))(via Possatto et al., 2011; Anastasopoulou et al., 2013; Foekema et al., 2013; Lusher et al., 2013; Di Beneditto & Awabdi, 2014; Neves et al., 2015; Phillips & Bonner, 2015; Bråte et al., 2016; Phillips & Bonner, 2016; Rochman et al., 2015; Rummel et al., 2016), demersal (15.5%; *n*=62, SD=24.0) (via Possatto et al., 2011; Dantas et al., 2012; Ramos et al., 2012; Anastasopoulou et al., 2013; Foekema et al., 2013; Lusher et al., 2013; Neves et al., 2015; Rochman et al., 2015; Miranda & de Carvalho-Souza, 2016; Phillips & Bonner, 2015; Phillips & Bonner, 2016; Rummel et al., 2016), pelagic (14.3%; *n*=53, SD=20.2), and bathydemersal (3.6%; *n*=7, SD=7.2) (via Anastasopoulou et al., 2013; Neves et al., 2015) zones. The bathydemersal category was severely limited in number of reported %FO values (*n*=7), and correspondingly fell well outside of the range of %FO exhibited by the other categories (all falling within the range of 14.3-16.3%) (via Anastasopoulou et al., 2013; Neves et al., 2015). The deviation of %FO within groups was high for all bathymetric zones ranging from 7.2% (demersal) to 29.8% (benthopelagic). The combination of “pelagic” with “bathypelagic”, and “demersal” with “bathydemersal” yielded no significant difference in the %FO between relative feeding zones (p=0.921, df=2, *n*=202)

##### The effect of species on ingestion rate

Some fish species were collected repeatedly in more than one study spanning different geographic regions. These species include: Atlantic horse mackerel (*Trachurus trachurus*) (Neves et al., 2015; Foekema et al., 2013; Lusher et al., 2013), john dory (*Zeus faber*) (Lusher et al., 2013; Neves et al., 2015), longnose lancetfish (*Alepisaurus ferox*) (Choy & Drazen, 2013; Jantz et al., 2013), small spotted catshark (*Scyliorhinus canicula*) (Neves et al., 2015; Anastasopoulou et al., 2013), and whiting (*Merlangius merlangus*) (Lusher et al., 2013; Foekema et al., 2013). The Atlantic horse mackerel, john dory, and longnose lancetfish had all ingested plastics with varying %FO, while the small spotted catshark and whiting had a mix of %FOs including 0%. The two sample groups of striped red mullet (*Mullus surmuletus*) were both collected from the Portuguese Coast, one wild, and one from fishmongers, and both had a 100% (%FO) (Neves et al., 2015). The blackbelly rosefish (*Helicolenus dactylopterus*) is the only species that was caught from two different marine regions and maintained a 0% (%FO) across both (Anastasopoulou et al., 2013; Neves et al., 2015). This included a large sample size of 380 individuals from the Ionian Sea (Anastasopoulou et al., 2013), and 1 individual from the Portuguese Coast (Neves et al., 2015). The effect of geographic location on ingestion rate

All fish, regardless of species, collected in the English Channel (Lusher et al., 2013), and in waters surrounding the Falkland Islands (Jackson et al., 2000), Hawaiian islands (Choy & Drazen, 2013; Jantz et al., 2013), and Mediterranean Sea (Anastasopoulou et al., 2013; Romeo et al., 2015), were found to have ingested plastic. By contrast, none of the species collected from the watershed flowing into the Gulf of Mexico had ingested plastic (0%FO for all) (Phillips & Bonner, 2015). The remaining geographic regions had a mixture of >%0 and 0% FO. The highest latitudes to the north exhibited 0% (%FO) while the frequency of occurrence in more centrally located regions was a mixture of 0% and >0%. Almost all of the studies looked at coastal regions, with a bias towards European marine landscapes (Table S1).

## Discussion

### Silver hake and plastic ingestion

This study is the first to investigate plastic ingestion rates in silver hake, and we report here a 0% frequency of occurrence for the species, a significant result for a species that has established and potential commercial value in human food economies. The vulnerability of a species to plastic ingestion (or lack thereof) may depend on several factors, including the geographical region that the species or population inhabits (Boerger et al., 2010; Foekema et al., 2013; Neves et al., 2015), the feeding behavior and ecology of the species in question (Moser and Lee, 1992; Peters et al., 2017; Silva-Cavalcanti et al., 2017), and the presence of plastics in the animal’s natural bathymetric feeding range (Dantas et al., 2012; Eriksson and Burton, 2003; Romeo et al., 2015).

### Geographical Region

The result of 0% for plastic ingestion in silver hake is unexpected on both counts. From a geographic perspective, the island of Newfoundland, and particularly the south shores, is not entirely removed from the influence of the North Atlantic Subtropical Gyre, an area where plastics are known to accumulate (Law et al., 2010). Moreover, the south coast of the island - where sampling occurred - is in proximity to the Gulf of St. Lawrence, an area that is subject to the introduction of plastics via numerous land and sea-based sources. The Gulf of St. Lawrence receives the output of the St. Lawrence River, a river servicing a watershed of approximately 100 million people (The St. Lawrence Seaway Management Corporation, 2016). The St. Lawrence River and its Gulf are also the home of an important shipping route, moving over 160 million tons of cargo per year (The St. Lawrence Seaway Management Corporation, 2016). Finally, Atlantic Canadian (inclusive of all provinces surrounding the Gulf of St. Lawrence) fisheries are responsible for over 84% of the national fishery by value and home to over 85% of the country’s registered fishing vessels (Fisheries and Oceans Statistical Services, 2016). Although no other English-language studies have investigated plastic ingestion rates of fish within the Gulf of St. Lawrence, a survey of surface plastics surrounding the North Atlantic Gyre verified their presence in the southern Gulf of St. Lawrence (Law et al., 2010). Here, silver hake are not ingesting marine plastics despite the presence of plastic within their regional waters.

At the same time, smaller regional scales matter. Previous studies have identified correlations between low plastic %FO and nearshore versus offshore collection sites (Anastasopoulou et al., 2013; Barnes et al., 2009; Browne et al., 2011, 2010; Collington et al., 2012; Davison and Asch, 2011; Dubaish and Liebezeit, 2013; Foekema et al., 2013; Liboiron et al., 2016; Lusher et al., 2015; Rummel et al., 2016). For example, Lusher et al.’s study (2013) that assessed the plastic contamination of 504 individual fish (10 species) sampled from North Atlantic coastal waters, found that 100% of the fish species collected within 10 km from shore (inshore) ingested plastics. Most notably were the following demersal fish: red gurnard (*Aspitrigla cuculus*; *n*=66), had a 51.5%FO for plastics, while 38% of Dragonets (*Callionymus lyra*; *n*=50) ingested plastics (Lusher et al., 2013). Contrastingly, Foekema et al.’s (2013) study in the North Sea included 1024 individual fish (6 species) caught offshore, where the plastic ingestion rate was significantly lower, at 2.6%. The geographic location of sampling in relation to the shoreline and relative proximity to land-based sources of plastic waste, then, appears to be an important parameter when examining the %FO’s of various species. As our silver hake were collected offshore (at least 62 km from the shoreline), then this may contribute to their ingestion rate, though it is doubtful it would result in a 0% rate for the entire sample.

### Food items and feeding habits

The type of food sought by different species is another variable that impacts the risk that a fish species will ingest plastics. Filter feeders, for example, such as herring and horse mackerel, have a higher chance of ingesting plastic than other fish species as food particles are indiscriminately filtered from surrounding water (Foekema et al., 2013; Rummel et al., 2016). In our literature review, these species had ingested a %FO of 2.5% (Foekema et al., 2013; Hermsen et al., 2017; Rummel et al., 2016) and a %FO of 18.3% respectively (Foekema et al., 2013; Lusher et al., 2013; Neves et al., 2015). Microplastics have been reported to resemble the potential prey of the animal, creating speculation that some species mistake microplastics for small prey (e.g. zooplankton) and directly target the plastics for ingestion (Anastasopoulou et al., 2013; Camedda et al., 2014; Ory et al., 2017). However, this theory is not applicable to higher trophic level predators, such as silver hake. Despite being opportunistic ambush predators (Helser and Alade, 2012), silver hake are known to be selective feeders (Bowman, 1975). All silver hake individuals (*n*=3) over the size of 40 cm in length had fed exclusively on fish, consistent with the conclusions of previous studies in that silver hake of 40 cm and larger tend to be exclusive piscivores (Durbin et al., 1983; Langton, 1982). The transition to exclusive piscivory begins when silver hake reach maturity (Vinogradov, 1984; Waldron, 1992). Prey analysis of silver hake in this study found that shrimp - a benthopelagic invertebrate - was the most common prey, followed by fish, in the digestive contents. This is consistent with the dietary transitioning described above, considering that the majority of individuals were under 40 cm in length but still mature, indicating they were in the process of transitioning to a piscivorous diet, hence the inclusion of fish in a diet otherwise dominated by shrimp (a smaller, more suitable prey item for individuals of a small body size). Regardless of the absence of an exclusively piscivorous diet in the individuals sampled here, the dominance of shrimp in the diet remains to be indicative of a highly selective feeding behavior. The predatory and selective feeding behavior of adult silver hake is likely to reduce their vulnerability to the primary ingestion of marine plastics, unlike generalist predatory fish which have been shown to have ingested relatively high rates (%FO) of marine plastics, such as king mackerel (62.5%) and sharpnose sharks (33%) off the coast of Brazil (Miranda and de Carvalho-Souza, 2016).

Despite the demersal categorization of silver hake (Froese and Pauly, 2017), the selective piscivorous feeding style of adult silver hake lends it to a predominantly pelagic feeding zone. Adult silver hake feed predominantly in the pelagic water column; beneath the surface where buoyant plastics accumulate and above the benthos where high density plastics accumulate. Fish at earlier life stages however, experience a more fluctuating diet from one largely based on fish in the spring and fall, to one including crustaceans and mollusks in the summer (Waldron, 1992). The high proportion of shrimp observed in the diet of fish analyzed by this study may correlate with the phytoplankton bloom and the corresponding growth of larval and juvenile shrimp on the Southern shelf of Newfoundland (Fuentes-Yaco et al., 2007). The phytoplankton bloom begins in mid-March, and peaks in May, which coincides with the time of year the silver hake were collected (Fuentes-Yaco et al., 2007). At the end of May, when the spring bloom subsides, larger juveniles will begin to transition to a more benthic, adult lifestyle, with the exception of males, which make vertical migrations to surface waters to feed at night (Fuentes-Yaco et al., 2007).

Secondary ingestion of plastics through the consumption of prey items containing ingested plastics provides a means of plastic ingestion regardless of the plastic’s density or the predator’s selectivity. Secondary ingestion is therefore likely to be a threat to silver hake. The absence of plastics recovered from silver hake in this study may indicate that the prey of silver hake are also not ingesting plastics frequently, although some common prey items come from families known to have ingested plastics (Koller et al., 2011; Waldron, 1992). It is also possible that the plastics ingested by their prey are < 1 mm and were not captured by our methods. This is possible considering that shrimp were found in 91% of fed fish and any plastics ingested by shrimp would likely be smaller than 1 mm in size.

### Bathymetric feeding range

The comparison of %FO values between bathymetric ranges in the global literature did not provide any significant insights towards why silver hake may not be ingesting plastics, neither when based on silver hake’s demersal classification, or its often pelagic feeding zone. The average %FO values for demersal and pelagic fish in the global literature were both near the top of the range, at 15.5% and 14.3%, respectively. Moreover, based on the existing data, the region of the water column that a species inhabits may not have an effect on the risk of plastic ingestion. These results are, however, far from conclusive, considering that most of the %FO values reviewed came from severely limited sample sizes, something that has the potential to skew results. The potential for small sample sizes to skew results is made evident in the range of average %FO values per bathymetric zone presented here. The average %FO per zone ranged from 3.6% to 16.3%, however, all zones clustered around the 14.3% to 16.3% with the exception of the bathydemersal category (3.6%). The bathydemersal category was limited by a small sample size (*n*=7) when compared to the other categories (*n*>35), likely a result of the technical difficulty of sampling demersal fish of the deep sea. This limitation in sample size may be to blame for the bathydemersal category representing such a strong outlier in the dataset, and the result of 3.6% may not be representative of bathydemersal fish as a whole.

While further research is required in order to determine the effect that bathymetric range may or may not have on the risk of plastic ingestion in fish, feeding behavior appears to heavily influence their %FO regardless of bathymetric range. For example, while the result of 0% for plastic ingestion in silver hake did not correspond with the average %FO of demersal fish (15.5%), it is in line with another fish that exhibits a similar feeding behavior. A total of 380 individuals of the blackbelly rosefish (*Helicolenus dactylopterus*) were analyzed by Anastasopoulou et al. (2013), and none were found to have ingested plastic. Both blackbelly rosefish and silver hake inhabit an overlapping geographic range and occupy similar depth ranges (Froese and Pauly, 2017). Despite their small difference in depth range, both fish are predators that transition from generalized feeding on invertebrates as juveniles to piscivorous, highly selective feeding as large adults (Figueiredo et al., 1995; Neves et al., 2011). The 0% (%FO) found in both fish may be a reflection of their selective feeding habits, a behavior that would result in the targeting of highly active prey (fish), reducing the likelihood that plastics would be mistaken for prey.

### The significance of 0%

In our literature review of ingestion studies, not a single paper reported a stand-alone rate of 0%. Yet many studies (*n*=11) contained species that had not ingested any plastics. We are concerned, as are others in the scientific community (Dickersin, 2005; Ekmekci, 2017; Granqvist, 2015; Hasenboehler et al., 2007) that studies that have found a 0% may not be published, or may only be published in instances where they are aggregated with other species so the overall rate is > 0%, resulting in a positive bias in publishing. Indeed, a new scientific journal, *New Negatives in Plant Science*, has recently been established specifically to reduce barriers for publishing negative and null findings. Indeed, the scarcity of published 0% plastic ingestion rates, paired with the fact that most 0% ingestion rates were reported for species in which only a small sample size was tested, severely limited our ability to detect trends in plastic ingestion based on bathymetric ranges across the global plastic ingestion literature. If 41% of ingestion rates are not reporting plastic ingestion, then this is a significant finding across species. Moreover, publication of more robust 0% rates allows us opportunities to re-focus our analyses towards behavioral and life history traits of individual species of fish‐‐ something that is common in plastic ingestion studies of other types of marine organisms, but less common for studies involving fish.

We conclude that even in the face of an extensive literature review of plastic ingestion reports across the globe, not enough data is yet available to make any broad-scale conclusions about trends in plastic ingestion based on bathymetrics, or even to characterize across species that have not been reported to ingest plastics. Further research might be conducted specifically on species that have reported 0% frequency of occurrence rates for plastic ingestion that have low sample sizes, first to understand whether these rates are due to low sample sizes or to the species generally, and secondly to better understand the behaviors, locations and other traits of fish that influence low or null percent ingestion rates. Understanding species that are less susceptible to plastic pollution is particularly important for places like Newfoundland, which depend on healthy fish for sustenance and economies.

## Contributions

Max Liboiron provided main ideation and structure for the article, co-wrote the article, mentored and trained all authors, and paid for the study.

France Liboiron conducted laboratory wet work, co-analyzed data, and co-wrote the article.

Justine Ammendolia contributed data for the literature review and co-wrote the article.

Jessica Melvin conducted the spectrometry, analyzed co-analyzed data, and co-wrote the article.

Jacquelyn Saturno co-wrote the article.

Alex Zahara co-wrote the article.

Natalie Richard collected samples.

## Acknowledgements

Research was made possible through a Marine Environmental Observation Prediction and Response Network (MEOPAR) grant, co-sponsored by Irving Ship Building (“Monitoring Marine Plastics in Canada’s North”). We would like to acknowledge Fisheries and Oceans Canada and the Fisheries and Marine Institute of Memorial University for providing support to obtain samples. We would like to thank Daigo Kamada for his help creating figures, Liang Zhu for manuscript revisions, the Merschod Lab for help with Raman Spectrometry, Wade Hiscock and Kiley Best for their help with sample collections, Susan Fudge and Laura Wheeland for coordinating the Celtic Explorer survey trip, and M. bilinearis for their personal sacrifices to the project.

We acknowledge that this science was conducted on the unceded, unsurrendered ancestral Lands of the Mi’kmaq and Beothuk. We would also like to acknowledge the Inuit of Nunatsiavut and NunatuKavut and the Innu of Nitassinan, and their ancestors, as the original peoples of Labrador.

---

**Table S1.**
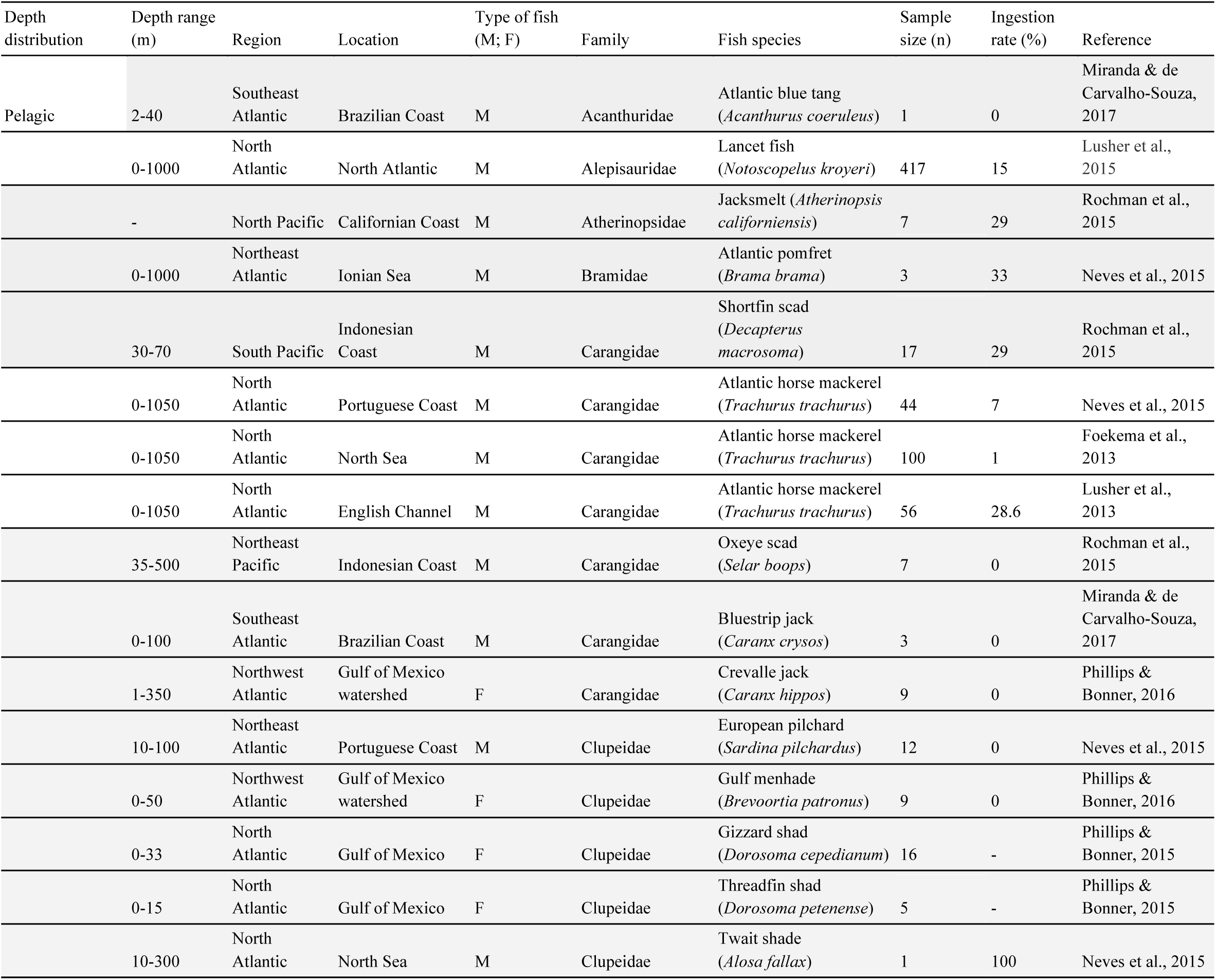
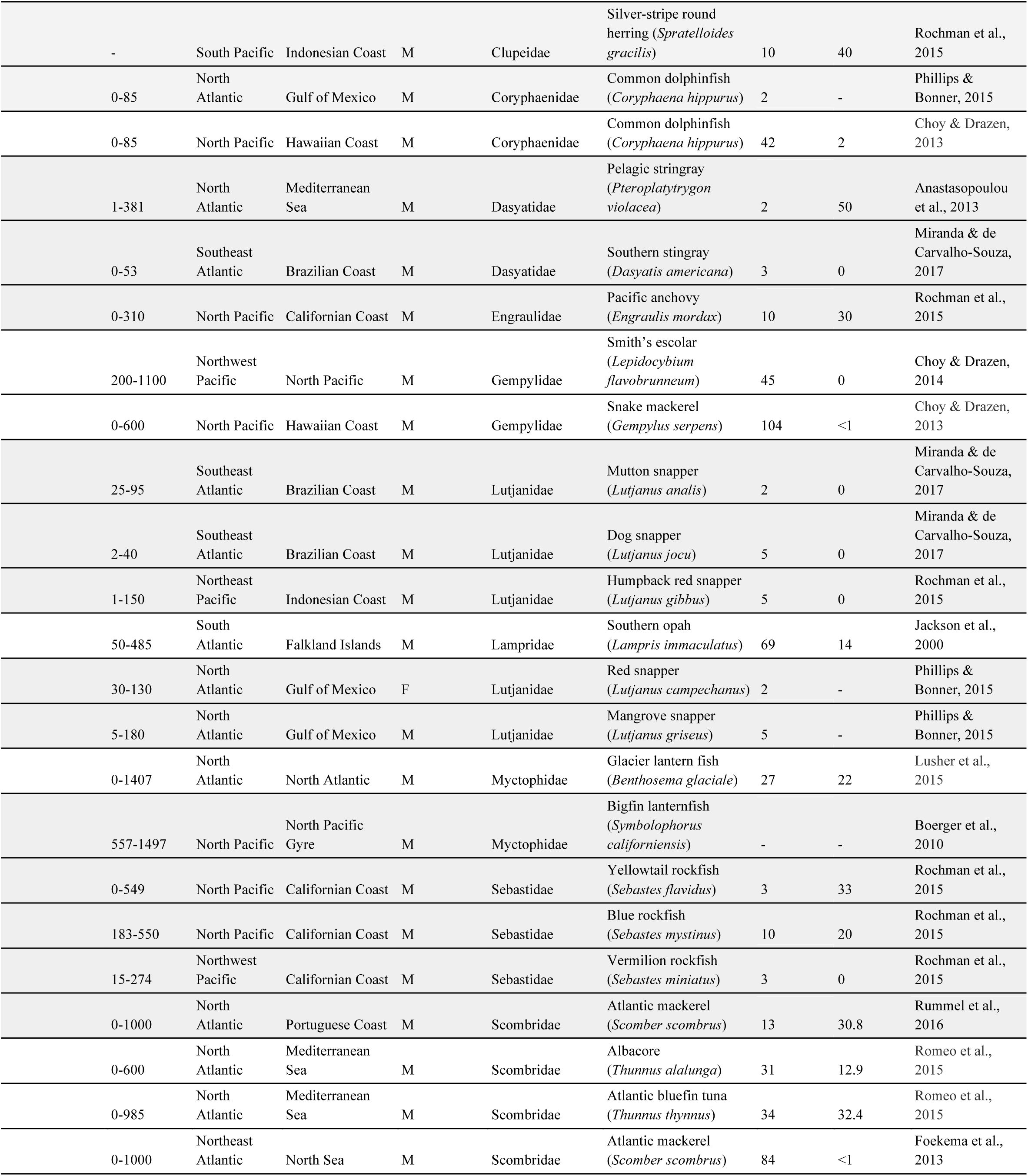
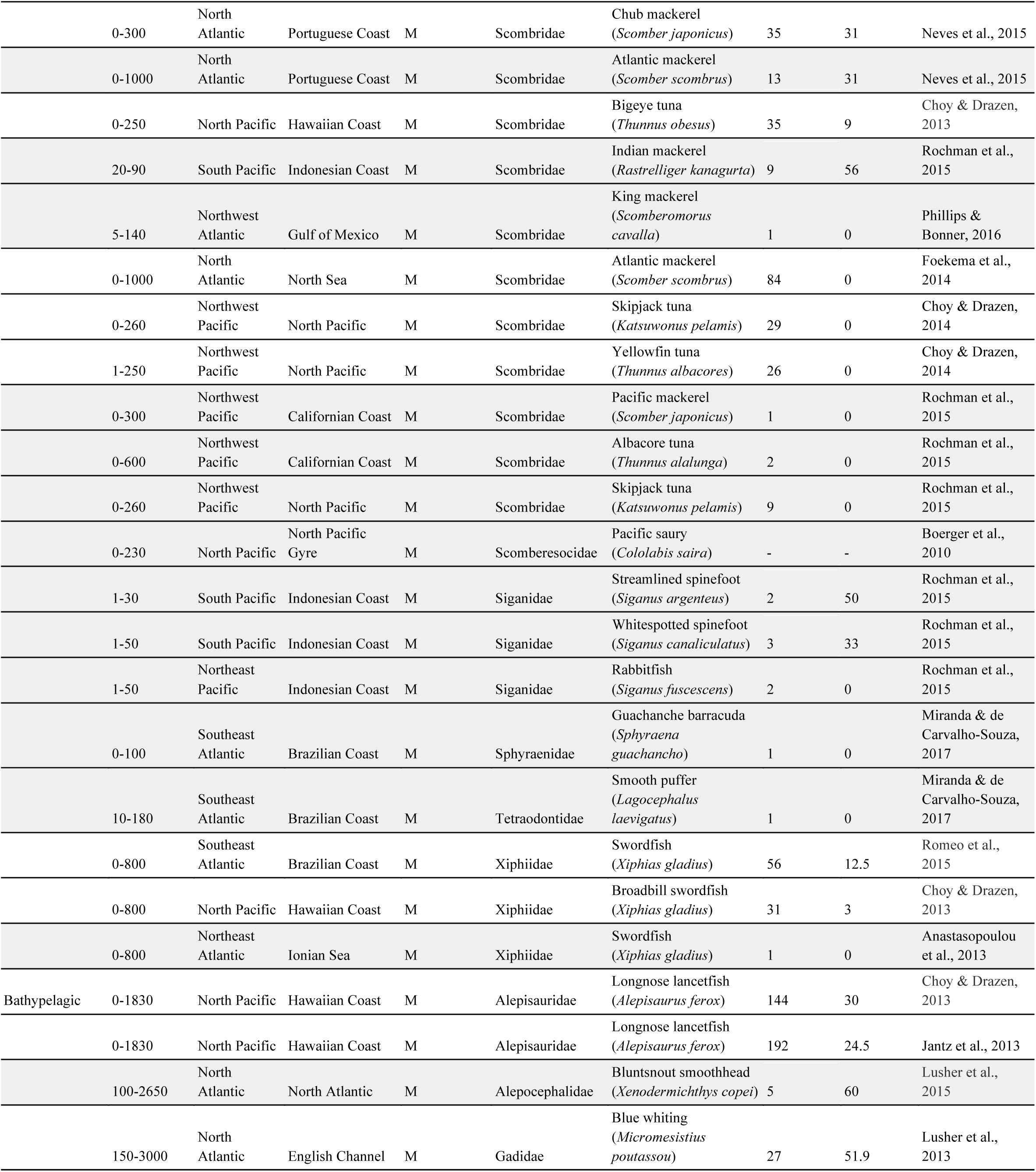
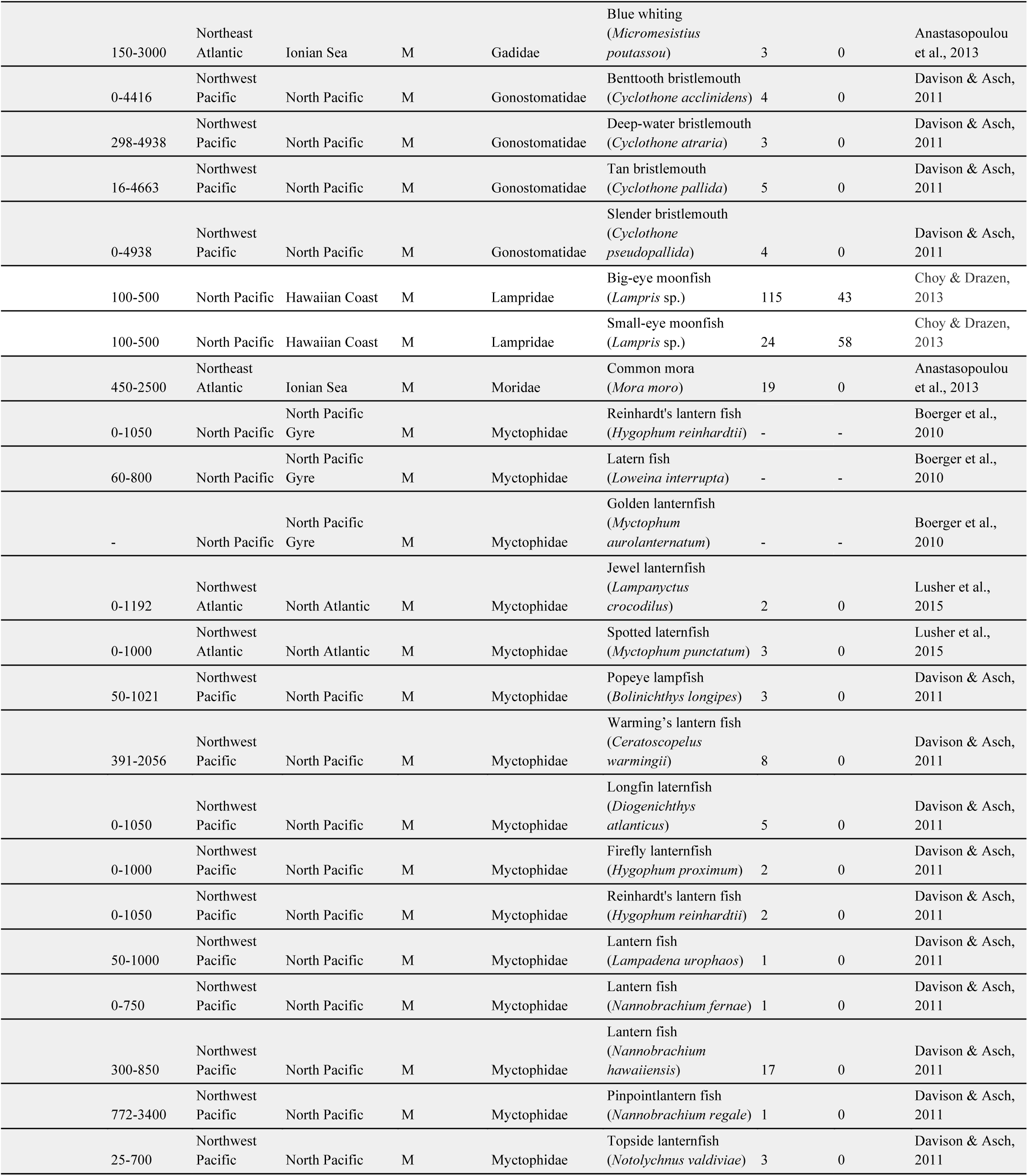
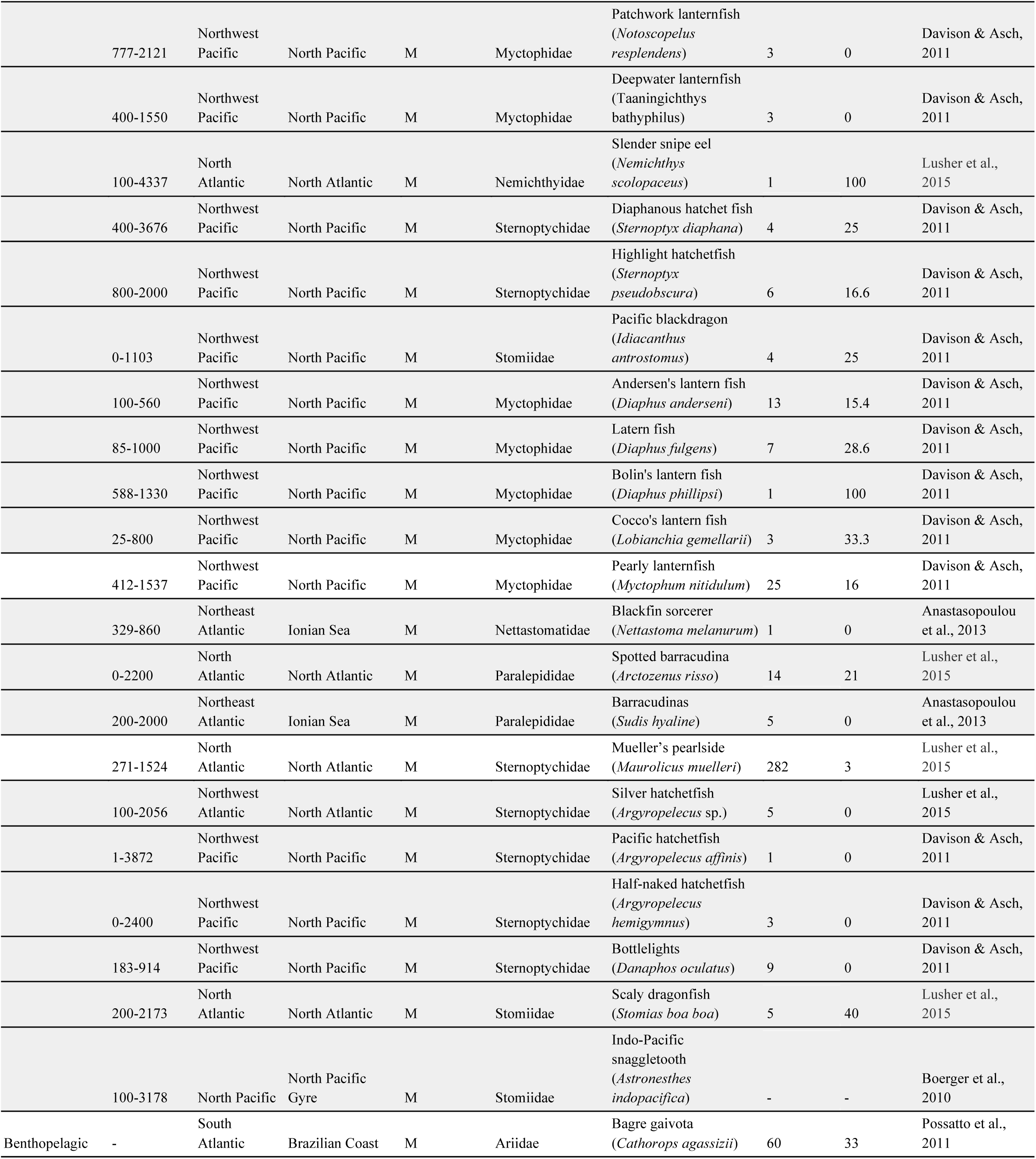
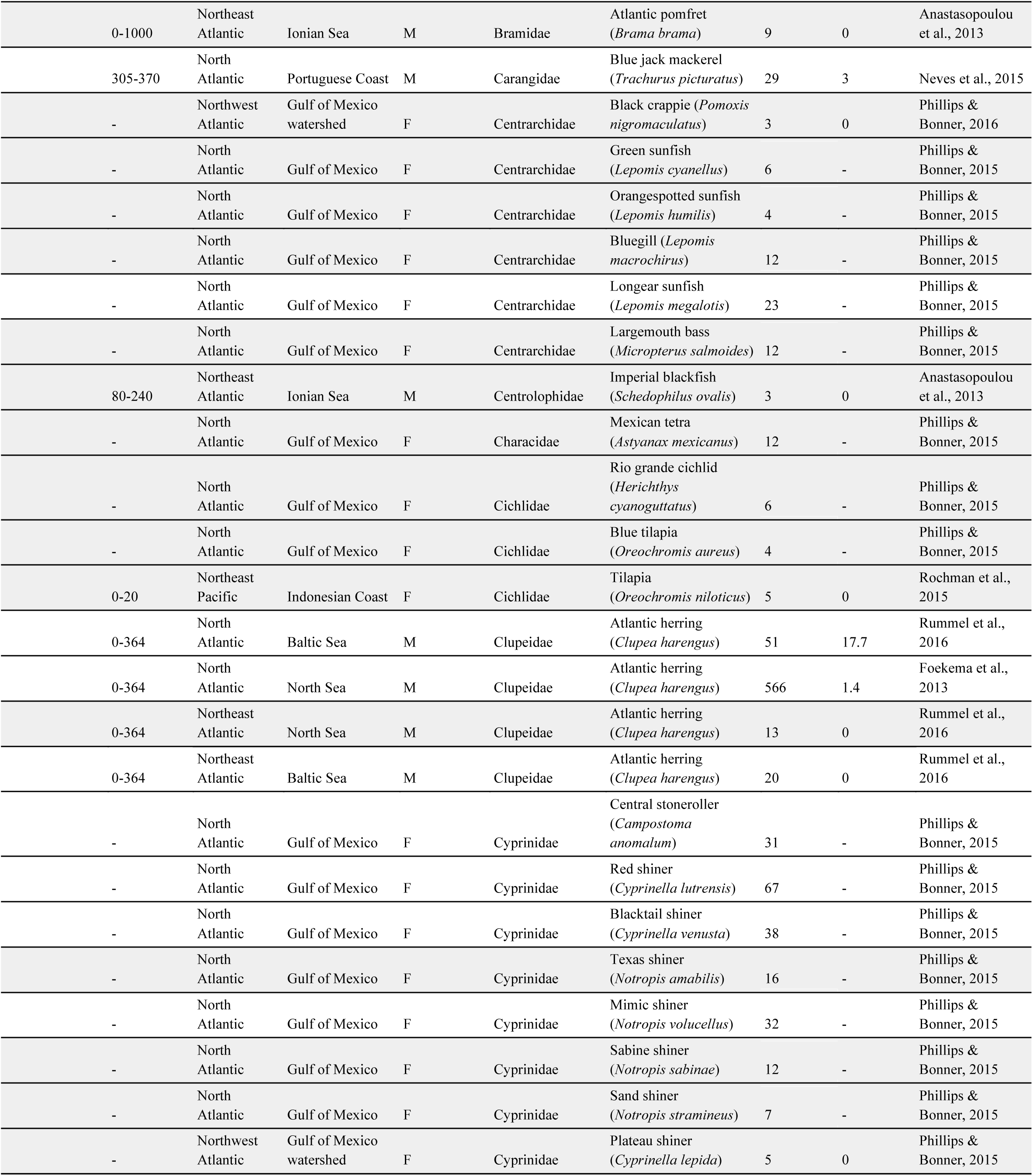
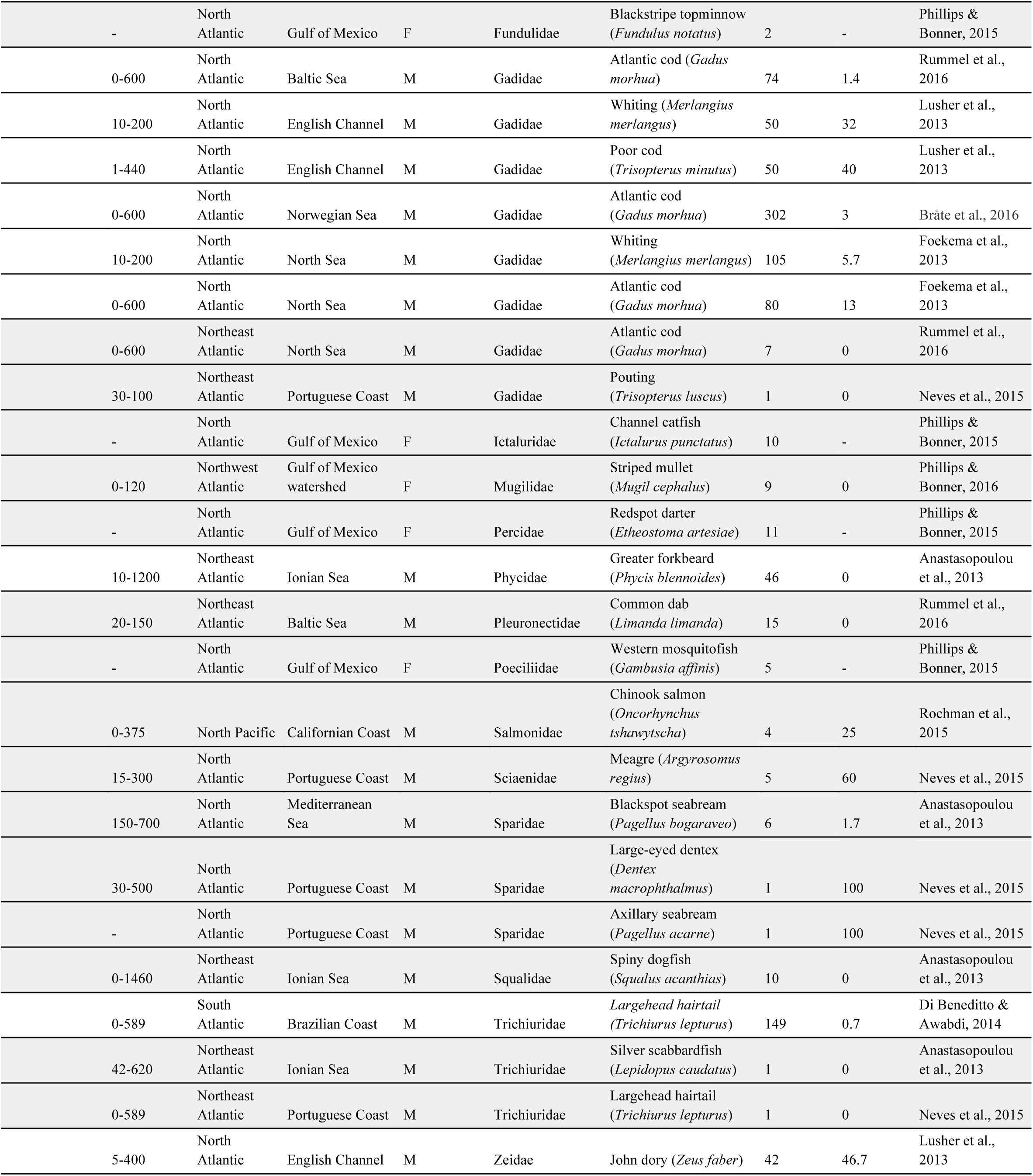
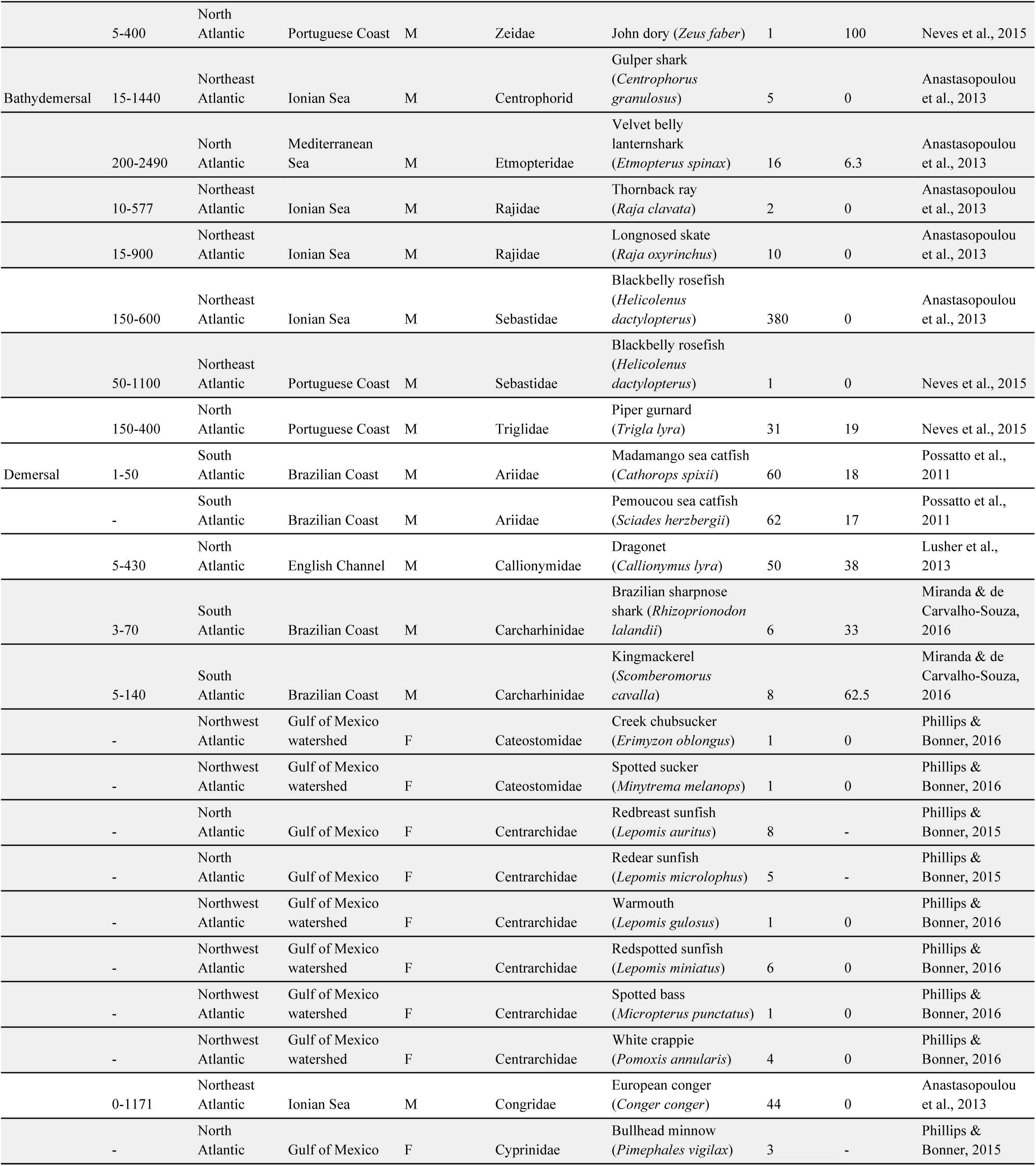
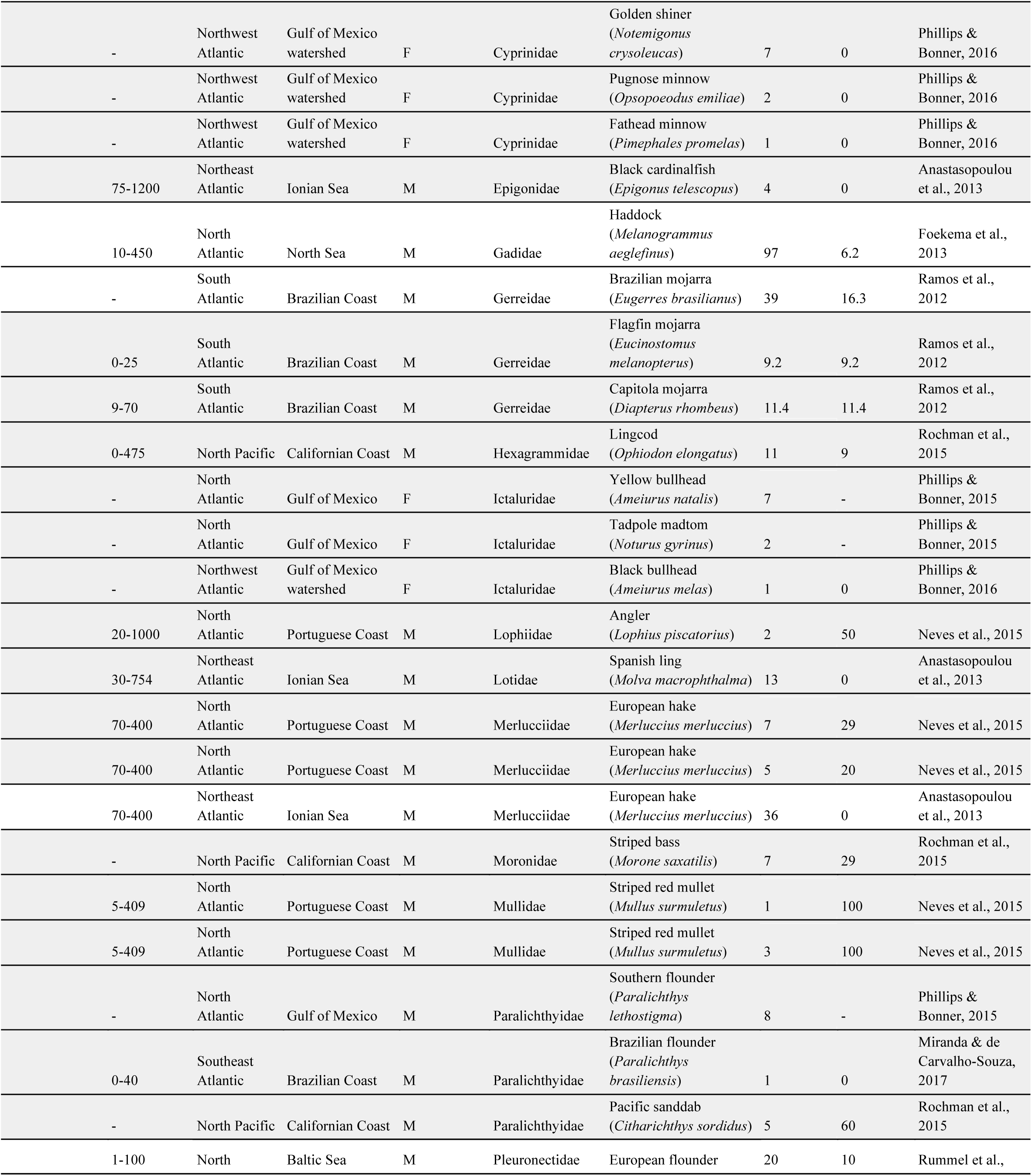
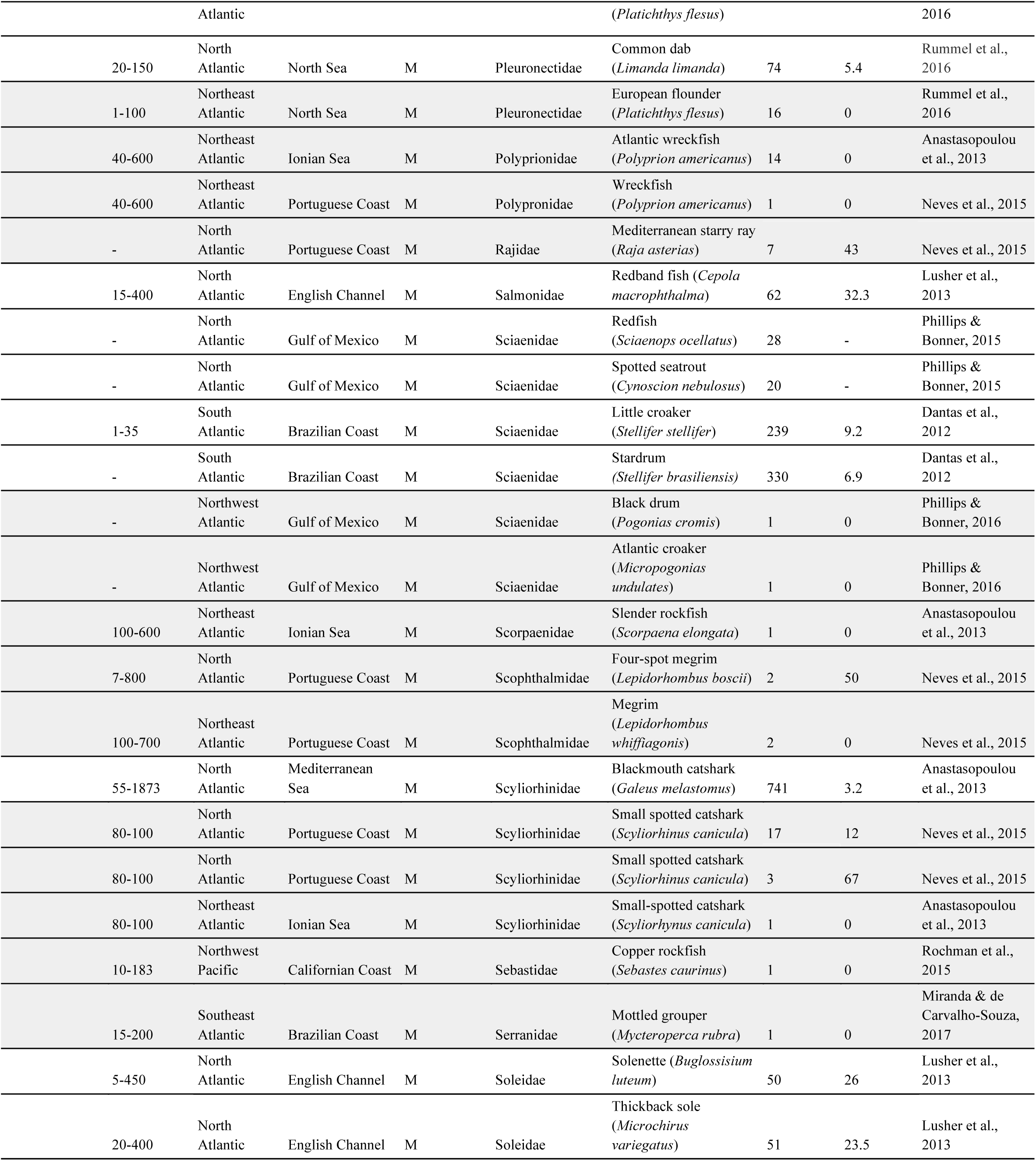
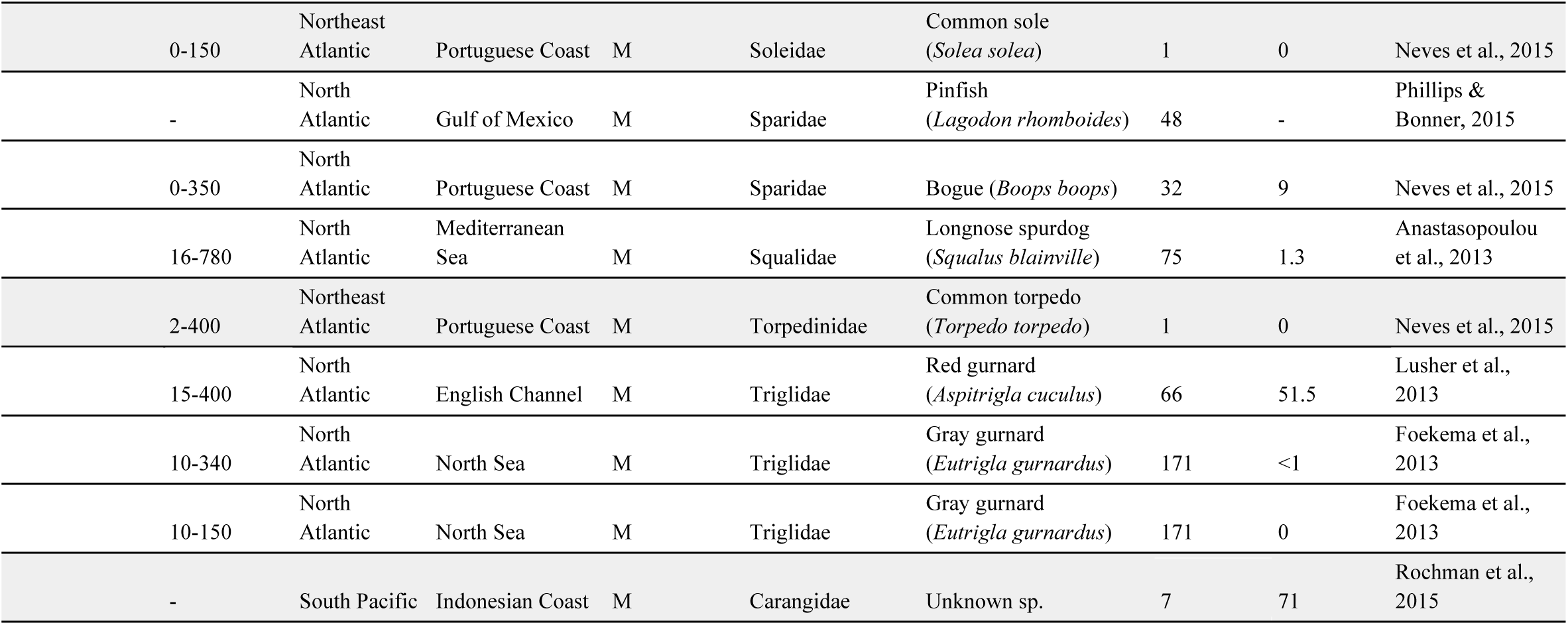
Marine plastic ingestion rates for both 0% and >0% in various species of fish from published studies. As some studies examined ingestion rates of multiple species, sample sizes broadly ranged from 1 to 741 individuals. Taxonomic information was provided to family and species level. This table was organized according to bathymetric depth distribution as determined by searching for species depth ranges and records on Fishbase (Froese and Pauly, 2015). Geographic region was categorized according to methods used by Liboiron et al., (2016) and specific location used included: Baltic Sea; Brazilian Coast; Californian Coast; English Channel; Gulf of Mexico; Gulf of Mexico watershed; Hawaiian Coast; Indonesian Coast; Ionian Sea; Mediterranean Sea; North Atlantic; North Pacific; North Pacific Gyre; North Sea; Norwegian Sea and; Portuguese Coast. The type of fish was defined by the habitat species occupy, marine (M) or freshwater (F). Grey shading of rows indicates that sample sizes sampled are less than 25 individuals whereas white shading represents that more than 25 individuals were sampled.

**Figure S1.**
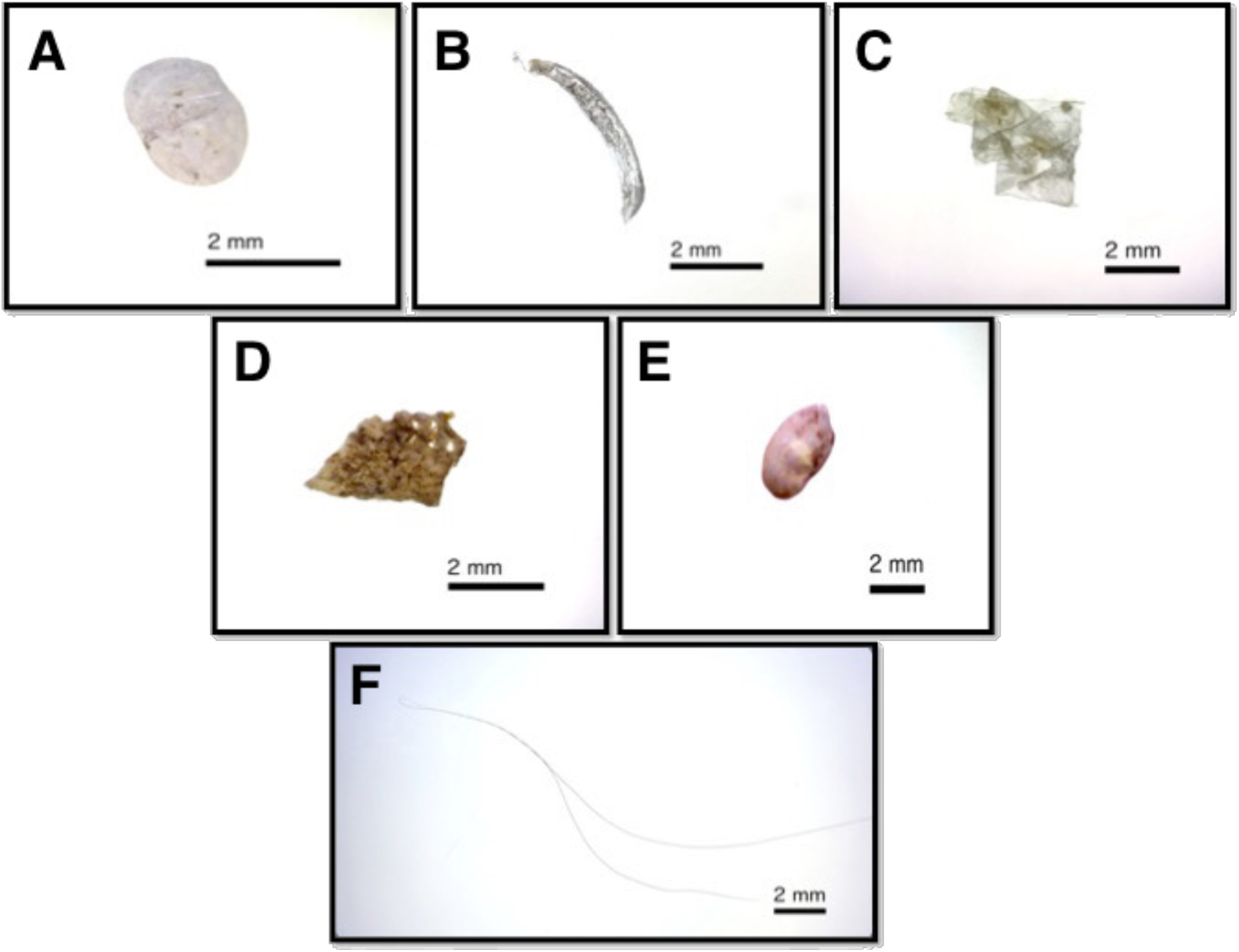
Particles obtained from the gastrointestinal (GI) tract dissections of silver hake individuals, and suspected to be of plastic composition. These particles were tested using Raman spectrometry and it was found that: (A-E) did not share similarity with the reference spectra for the common marine plastic polymers, but rather resembled fish bone fragments and, (F) was not logistically possible to process due to fine hair-like structure.

